# Metabolic function in aging retina and retinal pigment epithelium remains robust despite vision loss

**DOI:** 10.1101/2021.04.06.438698

**Authors:** Kristine A. Tsantilas, Whitney M. Cleghorn, Celia M. Bisbach, Jeremy A. Whitson, Daniel T. Hass, Brian M. Robbings, Martin Sadilek, Jonathan D. Linton, Austin M. Rountree, Ana P. Valencia, Mariya T. Sweetwyne, Matthew D. Campbell, Huiliang Zhang, Connor S.R. Jankowski, Ian R. Sweet, David J. Marcinek, Peter S. Rabinovitch, James B. Hurley

## Abstract

**Purpose:** Characterize how metabolic function in the murine retina and retinal pigment epithelium-choroid-sclera (eyecup) complex is impacted by natural aging.

**Methods:** We examined scotopic and photopic visual function of young (3-6 months) and aged (23-26 months) C57Bl/6J mice using electroretinograms (ERGs). Metabolic changes in retina and eyecup explants were characterized by measuring uptake and usage of U-^13^C-glucose or U-^13^C-glutamine at different timepoints by gas chromatography-mass spectrometry (GC-MS), measuring oxygen consumption rate (OCR) using a perifusion apparatus, and determining ATP levels with a bioluminescence assay.

**Results:** Scotopic and photopic ERG responses declined in aged mice. Glucose metabolism, glutamine metabolism, OCR, and ATP pools in retinal explants were mostly unaffected by the age of the mouse. In eyecups, glutamine usage in the Krebs Cycle decreased while glucose metabolism, OCR, and ATP pools remained stable.

**Conclusions:** The *ex vivo* approach in our study to examine aging glucose and glutamine metabolism in retina and RPE showed negligible impact of age on retina and an impairment of glutamine anaplerosis in eyecups. The surprising metabolic stability of these tissues *ex vivo* suggests age-related metabolic alterations in these tissues may not be intrinsic. Future experiments should focus on determining whether external factors including nutrient supply, oxygen availability, or other structural changes influence ocular metabolism *in vivo.*

## Introduction

Aging is associated with vision loss which contributes to disability and reduces quality of life^1,2^. Over the course of physiological aging, tissues in the eye exhibit functional decline. The retinal pigment epithelium (RPE) accumulates lipofuscin and drusen, Bruch’s membrane thickens, and overall cellular organization decreases^3–8^. These alterations are also characteristic of age-related macular degeneration, a severe disease associated with aging that is a leading cause of vision loss^2,9–12^.Within the neural retina, mitochondria are damaged^13–15^, synaptic connections between cells deteriorate^16^,^17^, and the functional capacity of photoreceptors is compromised.

In mice and humans, scotopic and photopic electroretinograms (ERGs) decline with age^18–21^. Manipulating metabolism *ex vivo* directly influences ERG response^22^. Increasing evidence suggests there is extensive metabolic interplay between the highly glycolytic retina and the retinal pigment epithelium^23–25^ and that metabolic changes in RPE can affect the retina. Increasing or disrupting glucose conduction in the RPE can result in retinal degeneration^26–28^. Similarly, aging represents a form of gradual, natural decay that impacts the eye. Metabolic changes are a hallmark of aging^29^ in many tissues. Although steady-state metabolite levels differ in tissues isolated from young and middle-aged mouse eyes^18^, to our knowledge the cross-talk between aging and central energy metabolism in the mammalian retinal ecosystem has not been examined. When characterizing age-related metabolic changes, it is not always clear whether the observed alterations are intrinsic to the tissue, or due to differences in vascularization or nutrient availability. An ex vivo approach to metabolic characterization^30^ can facilitate the identification of intrinsic metabolic changes.

To investigate the relationship between functional decline and energy metabolism in aging retina and RPE (Study structure in Figure 1), we measured visual function *in vivo* by ERG and interrogated intrinsic alterations of metabolism by analyzing *ex vivo* tissue explants using stable isotope tracers and targeted metabolomics.

**Figure 1:**
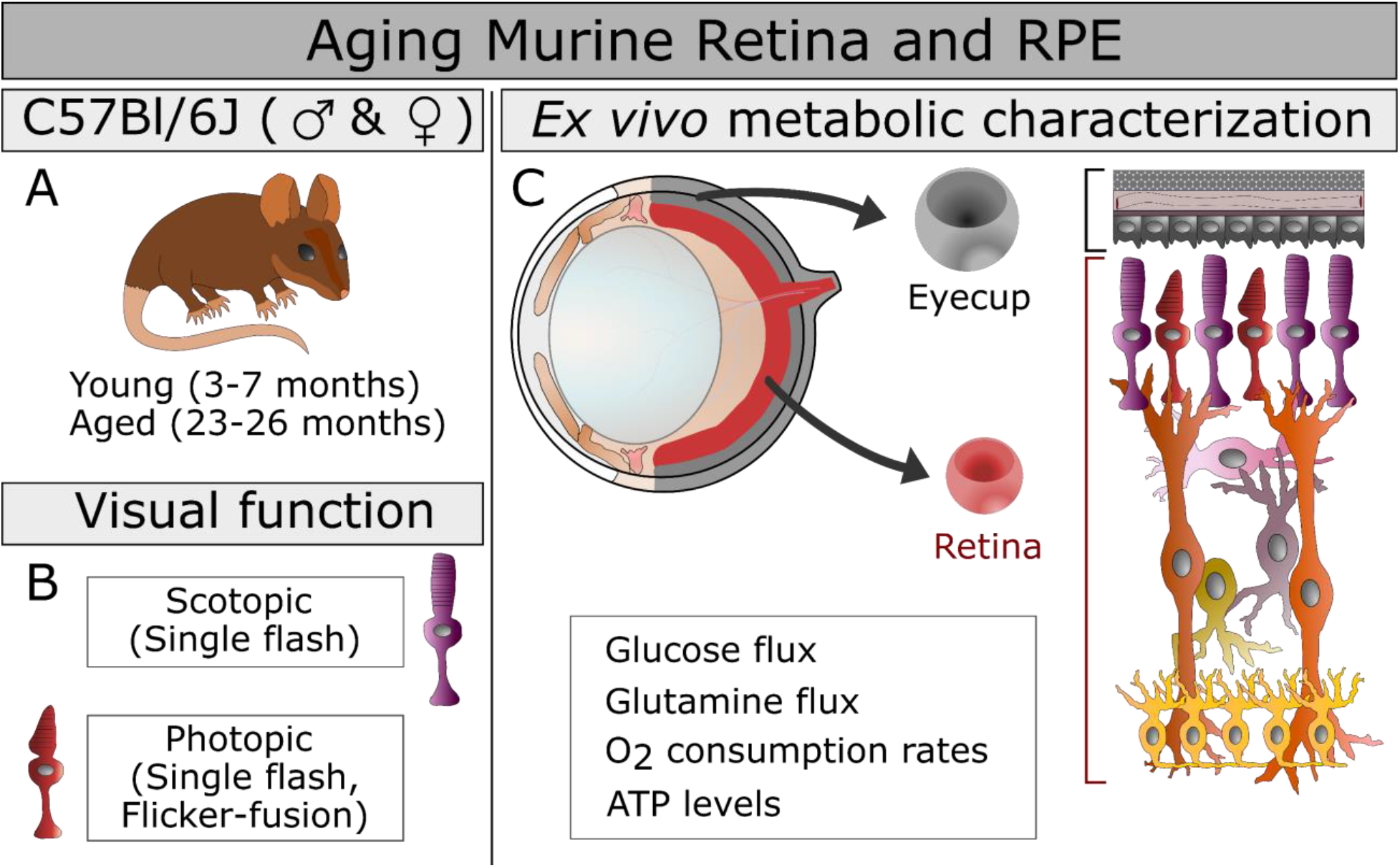
The study design is presented including the mouse groups, ages, and sex (A), the measures of visual function employed (B), and the *ex vivo* approaches to characterizing metabolism (C) in the aging eye. The functional metabolic measurements are listed below the structure of the murine eye (C, top left) from which we isolated two explants in this study (C, center): the retina RPE-choroid complex that includes the RPE, choroid, and sclera that has been cleared of connective tissue (Eyecup). The cellular composition of these explants (C, right) is highlighted with retinal cells shown in shades or red, purple, orange and yellow, while the eyecup is shaded in variations of gray.

## Methods

### Mouse ages, origin, and housing

C57Bl/6J mice of both sexes were obtained from the National Institute of Aging aged rodent colony (Bethesda, MD) and Jackson Laboratories (Bar Harbor, ME; Stock No: 000664). Young (3-7 months) and aged (23-26 months) mice were housed in groups of 5 or less in vivariums with *ad libitum* access to food (Rodent Diet 5053) and water. Some aged male mice were treated with saline via osmotic minipumps or provided water by bottle for 8 weeks prior to euthanasia as controls for separate studies unrelated to the eye. No differences were observed in ocular metabolism between untreated and control mice. Mice were confirmed free of the Crb1 mutation found in Rd8 models of retinal degeneration^31^,^32^. The light/dark cycle was 14 hours of light and 10 hours of dark. Experiments complied with the policies of the Animal Care and Use Committee at the University of Washington and the ARVO Statement for the Use of Animals in Ophthalmic and Vision Research.

### Electroretinogram (ERG) set-up

Mice were dark adapted overnight, anesthetized with isoflurane, and eyes dilated with 2.5 % phenylephrine (Akorn, Inc; NDC 174780201-15) and 1% tropicamide (Bauch + Lomb; NDC 24208-585-64). Gold electrodes were placed on each cornea. A reference and ground electrode were positioned on the back of the head. Mice were placed inside a UTAS Visual Diagnostic System with BigShot Ganzfeld with UBA-4200 amplifier (LKC Technologies; Gaithersburg, MD). Readings were taken from both eyes whenever possible, and the eye with the best response was analyzed.

### Single-flash ERGs

Recordings were elicited using flashes of LED white light at increasing flash intensities with two minute pauses between individual flashes under scotopic (−50 to 50 dB) and photopic (0 to 100 dB flashes, 30 cd/m^2^ background light) conditions. Readings were calibrated such that 0 db = 2.5 cd*s/m^2^. The a-wave amplitude was measured 8 ms after flash stimulus. The b-wave amplitude was measured as the magnitude from the a-wave minimum to the b-wave maximum.

### Flicker-fusion ERGs

Temporal resolution was measured using a 5 db flash of varied frequencies (20-50 Hz) under photopic conditions. Ten second pauses were taken between frequencies and repeated 9 times each.

Measurements were taken at 0.5 s intervals for a total of 512 points in 0.255 s (sampling frequency = 2003.91 samples/s). At each frequency, replicates were averaged. Figure 2E and 2F show examples of these raw values at 37 Hz. A waveform was generated and the magnitude calculated using Fast Fourier Transform (Microsoft Excel Data Analysis ToolPak). The sampling frequency was 2003.9 samples/s and the step value was 3.91.

**Figure 2:**
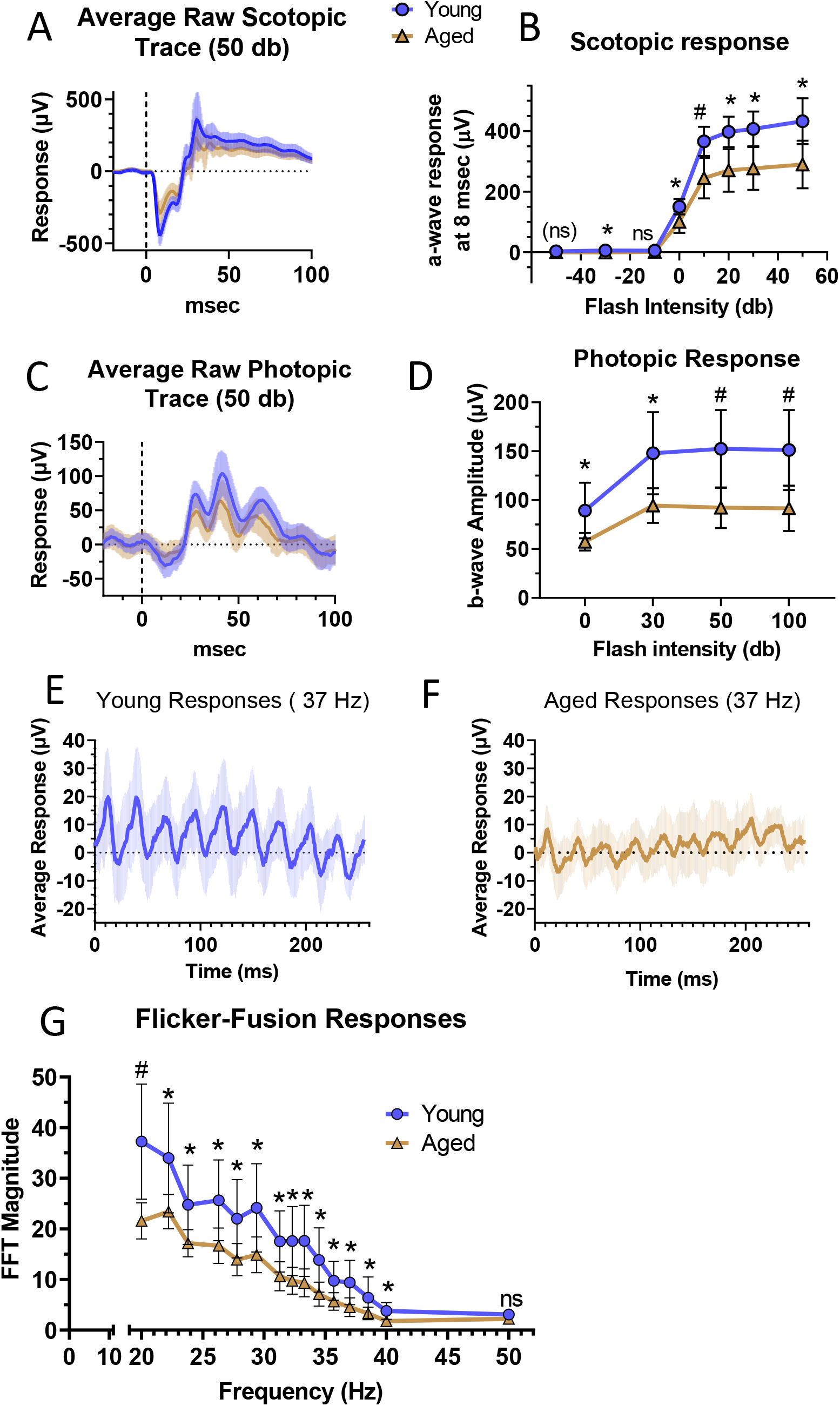
Single-flash ERGs were used to assess scotopic and photopic function in the retina. The averaged scotopic response curve in response to a 50 db flash (A) and the resulting a-wave amplitude from −50 to 50 db light at 8 ms after the flash (B) show a decline in the response of rod circuitry with aging. Sample sizes are 8 for young and 9 for aged. In photopic conditions (Background light of 30 cd/m^2^), the average response curve to a 50 db flash (C) and the resulting b-wave amplitude from flashes of 0 to 100 db (D) show a decline in the response of cone circuitry with aging. Sample sizes are 12 for young and 9 for aged. Photopic function and temporal resolution were examined with a 5 db flash which flickered between frequencies of 20-50 Hz. The averaged response to 5 db light flickering at 37 Hz is shown for young (E) and aged animals (F). The young have a more uniform and stronger response to equivalent stimuli than aged animals. Aging decreased the magnitude of the response at equivalent frequencies (G) as calculated by Fast Fourier Transform. Sample sizes are 8 for young and 9 for aged. Panels show the average ± standard deviation. Normality of data was determined using the Shapiro-Wilk test and p-values were calculated using unpaired t-tests (* = p < 0.05) or Mann-Whitney tests (# = p < 0.05, marker of non-significance enclosed in parentheses).

### Isolation of retinas and eyecups

Animals were euthanized by awake cervical dislocation. Eyes were enucleated, cleared of excess tissue, and the retina separated from the RPE-choroid-sclera complex (eyecup) in Hanks’ Balanced Salt Solution (Gibco; Grand Island, New York). Tissues for ATP determination were snap frozen and stored in liquid nitrogen. Those for flux or OCR were used immediately.

### Metabolic flux and metabolite extraction

Metabolite standards and buffer components were purchased from Sigma-Aldrich (MilliporeSigma; St. Louis, MO). Product numbers are included in Supplementary Table 2. Incubation medium was formulated as follows:

- Krebs’s Ringer Buffer (KRB): 98.5 mM NaCl, 4.9 mM KCl, 1.2 mM KH_2_PO_4_, 1.2 mM MgSO_4_, 20 mM HEPES, 2.6 mM CaCl, 25.9 mM NaHCO_3_
- Either 5 mM U-^13^C glucose alone or 5 mM unlabeled glucose and 2 mM U-^13^C glutamine U-^13^C metabolic tracers (99% isotopic purity) included ^13^C labeled D-Glucose and L-Glutamine (Cambridge Isotope Laboratories, Inc; Tewksbury, MA). Incubations, tissue extractions, and media extractions were performed as described previously^30^ with minimal changes. Some samples were split in half by volume before drying. All samples were spiked with methylsuccinate, L-norvaline, or L-norleucine as process and instrument controls. Metabolites were stored at −80°C until analysis.

### GC-MS analysis

Metabolite standards between 1.25-35 μM were used to generate calibration curves. The derivatization and selected ion monitoring (SIM) methods have been described in detail^30^ with few changes. The incubation length with N-tertbutyldimethylsilyl-N-methyltrifluoroacetamide (MilliporeSigma; St. Louis, MO) was increased to 60 minutes. Samples were analyzed using a 49 minute gradient on an Agilent 7890/5975C GC-MS system (Agilent Technologies; Santa Clara, CA). Two Agilent columns were used: a DB-5MS (128-5522) and an HP-5MS (19091S-433I) with flow rates of 0.8 mL/min or 1 mL/min, respectively. For all derivatized metabolites, we used target ions for quantification and isotopologue distribution determination, and a qualifier ion for identity confirmation. Supplementary Table 2 describes SIM method details for individual metabolites. Peak areas for SIM ions were obtained in MSD Chemstation (Agilent Technologies; Santa Clara, CA) with manual verification of automated peak integrations. IsoCor v2 was used^33,34^ to correct for natural ^13^C abundance and determine percent enrichment of ^13^C. In individual metabolite isotopologues, the number of incorporated ^13^C is represented shorthand by “Mx”, where x is the number of ^13^C (M0, M1, M2, etc.).

### Oxygen consumption rate

For each replicate, two retinas or four eyecups were quartered and loaded into a perifusion system that assesses oxygen consumption as described^35,36^ with minimal changes. Tissue was perifused with Krebs-Ringer buffer maintained at equilibrium with 21% O_2_, 5% CO_2_, and 74% N2 by an artificial lung and supplemented with 0.1 g/100mL BSA, 1X antibiotic-antimycotic (Gibco; Grand Island, NY), and 5 mM glucose. Succinate (5 mM) was added when testing mitochondrial function. Flow rate over live tissue averaged 61.9 ± 5 μL/minute. OCR was calculated as the product of flow rate times the difference in outflow and inflow oxygen levels. Data was reported as a change in OCR after subtracting off the baseline OCR measured in 5 mM glucose alone.

### ATP measurements

The pool of ATP in retinas and eyecups extracted in boiling water^37^ was measured via luminescence with the Molecular Probes^®^ ATP Determination Kit per the manufacturer’s instructions. Values were normalized to protein content in the pellet and supernatant.

### Protein concentration for normalization

Protein pellets were solubilized in RIPA buffer (150 mM NaCl, 1% Triton X-100, 0.5% sodium deoxycholate, 0.1% SDS, 50 mM Tris pH 8.0, 1X HALT protease/phosphatase inhibitor) and quantified using the Pierce™ BCA Protein Assay Kit per manufacturer’s instructions.

### Grouping and statistics

Sample collection from young and aged groups occurred at different times. Sample sizes represent biological replicates. Extractions, derivatization, and sample runs were processed in batches including both ages, genders, matched retinas, eyecups, and/or their media, and random timepoints. To account for circadian contributions, the time of death for animals used in glucose flux, glutamine flux, and OCR are plotted in Supplementary Figure 1. Normality of data was determined using the Shapiro-Wilk test. Differences between young and old animals were examined using unpaired t-tests (Indicated with ns or *) or Mann-Whitney tests (Indicated with (ns) or #), and those associated with age and sex were considered using Kruskal-Wallis and Dunn’s multiple comparison tests. Values that did not reach statistical significance are marked “ns”.

## Results

### Scotopic and photopic vision declines in aging mice

We recorded ERGs under scotopic and photopic conditions in young (4-5 months) and aged (26 months) male mice. Single flash scotopic ERGs measured rod-mediated vision. The amplitude of the a-wave was extracted from raw traces (Figure 2A) 8 msec after the flash and averaged (Figure 2B). Responses declined significantly between young and aged mice at −30 db and between 0-50 db. This is consistent with previous findings that a-waves decline in mice as early as 16 months^18^,^19^.

Cone-mediated vision was characterized by examining the b-wave amplitude in raw traces (Figure 2C) in single-flash photopic ERGs. The b-wave amplitude declined significantly in aged animals relative to young (Figure 2D) at the flash intensities tested. Temporal resolution was considered by flicker-fusion ERG, which used a 5 db flashing light at frequencies between 20-50 Hz. An overall loss in temporal resolution was seen in the raw responses. Examples of averaged responses at 37 Hz are shown from young (Supplementary Figure 1A) and aged (Supplementary Figure 1B) mice. After Fast Fourier Transformation, the response magnitude was found to decline in aging (Figure 2E, main panel). Although the degree of change was variable, the magnitude declined significantly with age at all frequencies tested except 50 Hz.

### Metabolic flux from glucose is preserved in retinal explants cultured *ex vivo*

We sought to determine if glucose usage in the retina and RPE were altered at an age with known functional decline (Figure 2). Retina and eyecups were isolated from young (3-5 months) and aged male (26 months) and female (23-25 months) mice. Explants were incubated in KRB with U-^13^C-Glucose for 2, 10, 20, 30, or 45 minutes. Labeled metabolites generated from glucose in tissues and incubation media were measured by GC-MS (Figure 3A). Metabolites were quantified in terms of pmol or nmol metabolite per μg of protein. We examined metabolites in the tissue or exported by the tissue and considered the percent of the total pool labeled with ^13^C, and product-reactant ratios for particular steps in glycolysis and the TCA cycle.

**Figure 3:**
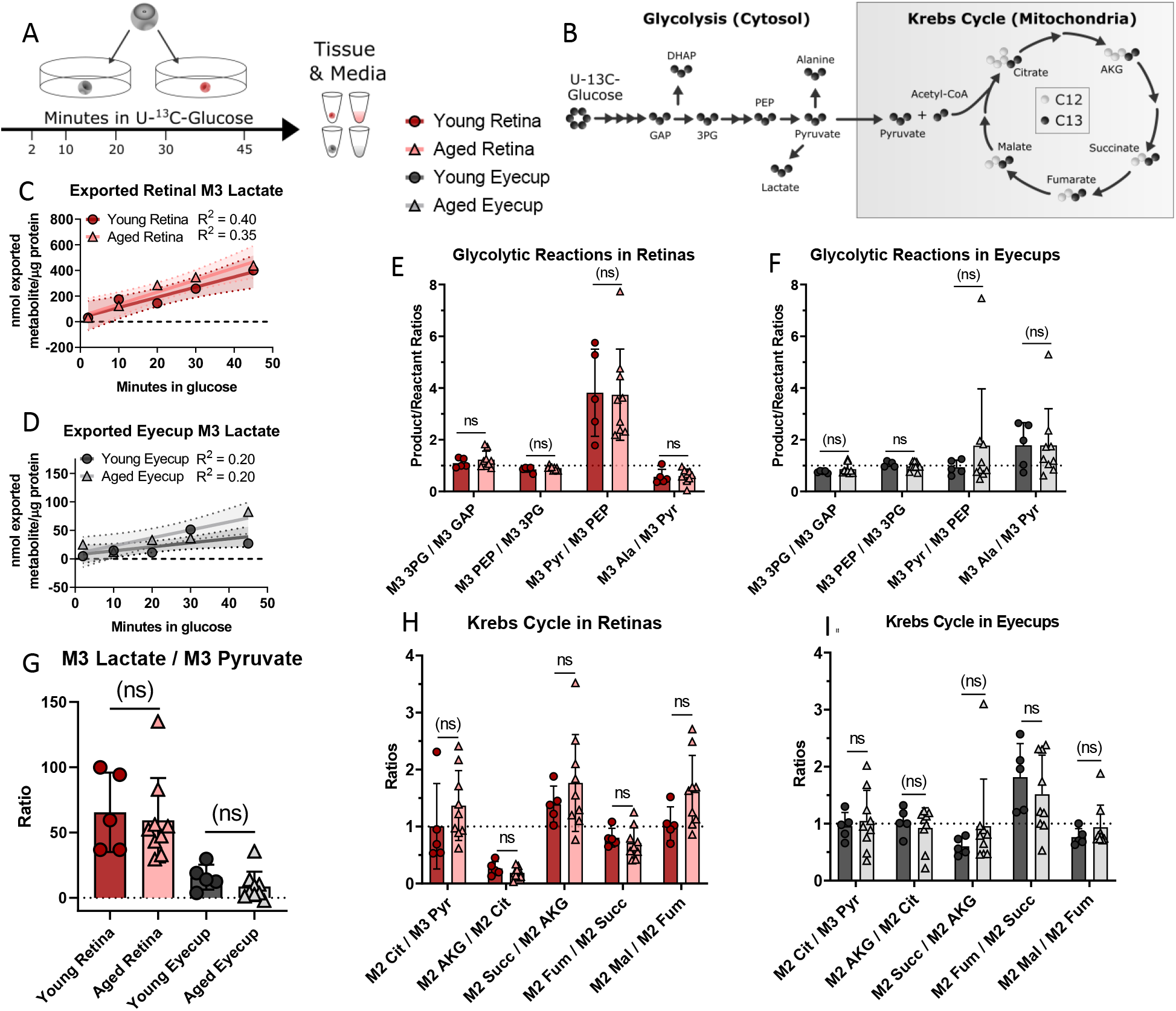
Metabolic activity was examined by incubating retinal and RPE-choroid (eyecup) explants in U-^13^C-glucose between 2-45 minutes. The tissue was washed and frozen, and aliquots of the incubation media were collected for analysis (A). Labeled intermediates downstream of glucose were quantified (B) in terms of percent ^13^C incorporation, pmol of ^13^C-labeled isotopologue per μg of protein in the retinal or eyecup explant, and product:reactant ratios for glycolytic reactions and those pathways that can be entered via pyruvate. Percent incorporation, pool sizes, and amount of isotopologue are shown in Supplemental Figure 3 and 4. To examine glycolytic activity, the amount of exported M3 lactate was measured in the incubation media of retinas (C) and eyecups (D). The slope of the lines were all non-zero in retina (p_young_ = 0.002, p_aged_ = 0.0003) and eyecups (pyo∪⊓g = 0.03, p_aged_ = 0.009), but showed no significant age-related change (Regression slope: pretina = 0.7, peyecup = 0.3). Within tissues, product:reactant ratios in glycolysis and common exit points to other pathways were plotted at 30 minutes because both tissues had reached a steady state. No significant age-related changes were seen in retina (E, G) or eyecup (F, G). Moving into the Krebs Cycle (B) at 30 minutes, no statistically significant age-related were observed in retina (H) or eyecup (I) explants. Normality of data was determined using the Shapiro-Wilk test and p-values were calculated for age-related comparisons using unpaired t-tests (* = p < 0.05) or Mann-Whitney tests (# = p < 0.05, marker of non-significance enclosed in parentheses). Error bars represent the standard deviation, except in Panels C and D which show the 95% confidence interval for the linear regression. **Abbreviations:** Glyceraldehyde 3-phosphate (GAP), Dihydroxyacetone phosphate (DHAP), 3-phosphoglycerate (3PG), Phosphoenolpyruvate (PEP), Pyruvate (Pyr), Alanine (Ala), Citrate (Cit), α-ketoglutarate (AKG), Succinate (Succ), Fumarate (Fum), Malate (Mal)

In tissues, we first examined glycolytic intermediates (Figure 3B) in retina and eyecup. We found no consistent change in pool size, percent ^13^C-incorporation, and labeled isotopologues in either tissue (Supplementary Figure 2). There was no significant change in export of M3 lactate from retinas (Figure 3C) or eyecups (Figure 3D) into incubation medium. We examined product:reactant ratios at 30 minutes in both tissues when label incorporation has stabilized. There were no substantial age-associated changes in glycolytic reactions in the retina (Figure 3E and 3G) or eyecup (Figure 3F and 3G). Glycolytic function appears preserved in both tissues.

We next considered how age impacts the Krebs cycle. As with glycolysis, we saw no consistent changes in retinas (Supplementary Figure 3A-C) or eyecups (Supplementary Figure 3D-F) in pool size, percent ^13^C incorporation, or labeled isotopologues. There were no significant age-related changes in product:reactant ratios in retinas (Figure 3H) or eyecups (Figure 3I) at the same 30-minute time point considered for glycolysis. Age did not significantly impact the Krebs cycle in retinas and eyecup explants.

Sexual dimorphism can influence outcomes in biological studies^38^, and can differentially impact the visual system generally^39^ and in the context of aging^40,41^. We considered glucose processing in young and aged female mice in parallel to males at 2 minutes. Total protein content in retinas was unchanged between males and females (Supplementary Figure 1A), however in eyecups (Supplementary Figure 1B) there was a change in aged males that was accounted for by normalizing all values to protein content in tissues. There were no significant contributions of age or sex in pool size or isotopologues (Supplementary Figure 4A and B) in retina or eyecups (Supplementary Figure 4C and D). No sex-dependent differences in aging retinal or eyecup metabolism were found.

### Glutamine metabolism is altered *ex vivo* in aged eyecups

Mitochondrial dysfunction is a known hallmark of aging^29^. To more directly investigate mitochondrial metabolism in aged ocular tissues we examined the usage of glutamine and succinate, which are oxidized by mitochondria in the RPE^42,43^.

Glutamine usage in *ex vivo* eyecup and retina explants was measured in 5 mM ^12^C-glucose and 2 mM U-^13^C-glutamine for 20 or 90 minutes. Metabolite pool size was diminished in aged eyecups at both timepoints (Supplementary Figure 5A) in glutamine, glutamate, and the downstream Krebs cycle metabolites α-ketoglutarate, fumarate, malate, and aspartate. Since we did not observe this defect in eyecups supplied with glucose alone (Figure 3), we hypothesized aged eyecups may have a defect in mitochondrial glutamine metabolism. After incubation in ^13^C labeled glutamine, M5 glutamine and glutamate (Figure 4B and 4C) were not significantly lower in aged eyecups. However, levels of M5 α-ketoglutarate, M4 Fumarate, M4 malate, and M4 aspartate (Figure 4D-G) did trend lower in aged eyecups compared to young. The percent labeled with ^13^C (Figure 4H-M) varied by metabolite. Notably, M5 glutamine, M4 fumarate, M4 malate, and M4 aspartate were consistently lower in young eyecups at 20 minutes, but had essentially matched the aged by 90 minutes. No significant changes in product:reactant ratios (Supplementary Figure 5B) occurred with age. These data suggest that glutamine anaplerosis is diminished in the aged eyecup.

**Figure 4:**
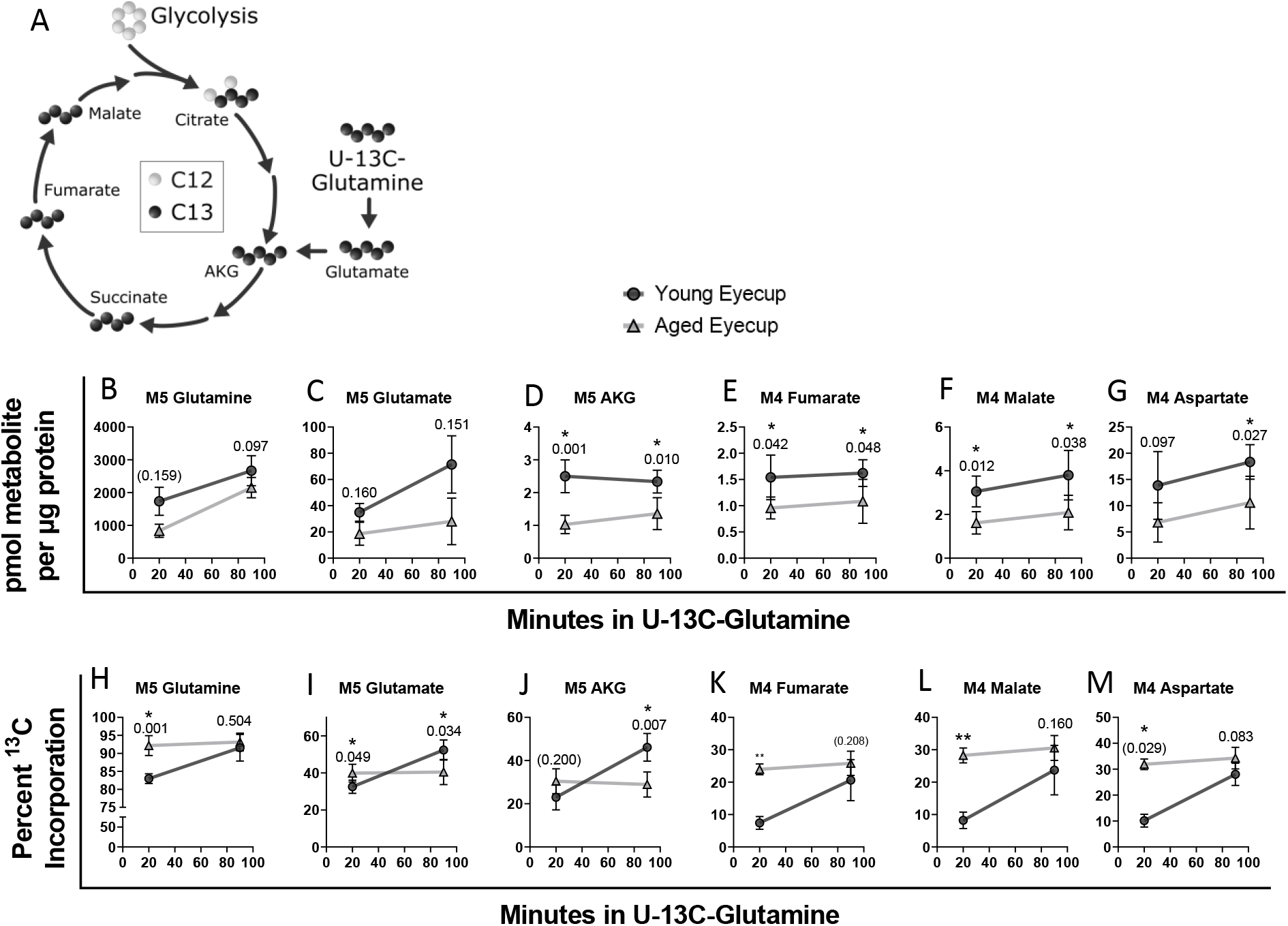
Metabolic activity with glutamine - and in the mitochondria more broadly - was examined by incubating eyecup explants in U-^13^C-glutamine for 20 or 90 minutes. The M4 and M5 labeled isotopologues entering into and proceeding through the Krebs cycle (A) were quantified in terms of pmol of ^13^C-labeled isotopologue per μg of protein in eyecup explants (B-G). Glutamine (B) and glutamate (C) trended lower in the aged eyecups at both timepoints examined, but did not reach statistical significance. In the downstream intermediates AKG (D), fumarate (E), malate (F), and aspartate (G) that decline was increased. The percent incorporation of 13C remains essentially unchanged in aged eyecups at both timepoints (H-M). Percent ^13^C incorporation rises past the aged in glutamate (I) and AKG (J). However, in glutamine (H), fumarate (K), malate (L), and aspartate (M), the percent ^13^C incorporation is consistently lower in young eyecups at 20 minutes, but had essentially matched the aged by 90 minutes. These changes can be related back to pool size and product:reactant ratios, which are included in Supplemental Figure 6. Normality of data was determined using the Shapiro-Wilk test and p-values were calculated (* = p < 0.05, ** = p < 0.0001) for age-related comparisons using unpaired t-tests or Mann-Whitney tests (# = p < 0.05, marker of non-significance enclosed in parentheses). Error bars represent the standard deviation.

The same analysis was performed in retinas. In contrast, no significant age-related changes in pool size or ^13^C-labeled intermediates (Supplementary Figure 6A and C) were observed. The percent incorporation at 20 minutes was higher in later Krebs cycle intermediates but remained unchanged by 90 minutes while younger retina label uptake trended higher (Supplementary Figure 6B). No significant changes in product:reactant ratios (Supplementary Figure 6D) were observed.

We postulated the declines in eyecup glutamine metabolism indicated a decline in mitochondrial function. We approximated mitochondrial function by measuring oxygen consumption rate (OCR) and ATP levels in young and old retinas and eyecups. Succinate is a mitochondrial respiratory substrate that is preferentially oxidized by the RPE^42^. Succinate-driven OCR was calculated for each animal by taking the maximum steady-state OCR of tissue respiring on 5 mM glucose + 5 mM succinate and subtracting the 5 mM glucose baseline OCR (Supplementary Figure 7A) and retinas (Supplementary Figure 7B). Steady-state OCR values are listed in Supplementary Table 1. As reported previously, succinate stimulates OCR in eyecups to a significantly greater degree compared to retina^42^. Neither retinas nor eyecups exhibited significant age-related changes in succinate-stimulated O_2_ consumption (Supplementary Figure 7C, 8D). Steady-state levels of ATP (Supplementary Figure 7E) also showed negligible changes.

## Discussion

Aging drastically alters visual function in humans^20,21^. We observed the same trend in 26 month-old C57Bl/6J mice where rod function, cone function, and photopic temporal resolution declined relative to young mice (Figure 2). By 26 months, C57Bl/6J mice are roughly equivalent to 79 year-old humans^44^ and functional changes are representative of advanced age, as roughly 25% of the population would have died^45^,^46^.

Rapid aerobic glycolysis in the retina is required for visual function^22,24^. Despite the observed visual decline in aged mice, the metabolic function of the retina in terms of glucose and glutamine metabolism is stable *ex vivo.* Glycolysis had reached a steady-state, but both glycolytic and Krebs cycle intermediates (Figure 3, Supplementary Figure 2, 3) were essentially unchanged between young and aged retinas. There was no observed effect of sex (Supplementary Figure 4), and mitochondrial OCR and ATP levels were also unchanged. Glutamine metabolism is linked to synaptic transmission through the neurotransmitter glutamate^47^. However, it is unlikely the minor defect we observed in the percent incorporation of ^13^C from U-^13^C-glutamine in late Krebs cycle intermediates (Supplementary Figure 6) will impact neurotransmission. We postulate that aging retina is robust and retains metabolic function on par with young retinas despite significant physiological decline *in vivo.*

Glucose metabolism appears to also be preserved in aging eyecups (Figure 3, Supplementary Figures 3 and 4). However, glutamine metabolism in aged eyecups is significantly different from young. The pool size (Supplementary Figure 5) and isotopologues were decreased with age. However, the percent ^13^C incorporation into glutamine intermediates (Figure 4) remained stable in aging eyecups compared to the increasing percent in young. We hypothesize that this indicates aged eyecups are deficient in glutamine metabolism and reach their maximum metabolic capacity faster than their younger counterparts. While the decreased M5 Glutamate:M5 Glutamine ratio could be indicative of decreased glutaminase activity in aging RPE, enzyme activity assays from glutaminase to fumarase would allow more targeted identification of the source of the defect in glutamine metabolism we observed.

Both metabolic changes and mitochondrial dysfunction are hallmarks of aging^29^. Glycolytic capacity declines with age in tissues like the brain and effector T Cells^48–51^. Young hearts subsist on fatty acid oxidation, but switch to glycolysis and ketone body usage in aging and heart failure^52,53^. Liver and skeletal muscle experience metabolic shifts with age^54–59^. Our results show that metabolism in retina and RPE are surprisingly robust in aging relative to these other systems despite functional visual decline in aged mice *in vivo.* This may indicate that the main driver of metabolic change in the aging eye is not intrinsic, but is due to nutrient or oxygen availability from blood. Blood vessel function around the retina has been shown to decline in aging people^60–62^. Oxygen extraction is lower in aged human retina^63^. Retina and eyecup explants are exposed to ambient oxygen and levels of glucose (5 mM) that are physiological to young mice, with few barriers to transport. Within the aged animal, these tissues may have restricted access to nutrients and oxygen due to the aforementioned declines in circulation and oxygen extraction. While retina and eyecup maintain their metabolic function in aging *ex vivo,* they may still experience a non-cell autonomous functional decline.

### Considerations for future studies

A complementary *in* vivo approach is necessary to determine whether metabolic changes in the aging eye are due solely to intrinsic properties of the tissues or may be related to alterations in the vasculature and tissue structure. We will utilize direct infusion of ^13^C-labeled fuels^64^ to test this hypothesis. Our study focused on fuels of known importance in retina and RPE (glucose, glutamine, and succinate), but other metabolic pathways known to be important in these tissues merit consideration in the aging eye^42,43,65–68^.

Finally, it is important to consider that mitochondria have diverse functions outside of cellular respiration^69–71^. Abundance of Krebs cycle intermediates provides a direct link to epigenetic change^72^,^73^. Mitochondrial dysfunction is also involved in aging and cellular signaling that can have broad implications in an organ system^29^,^70^,^74–77^. While the tissues in the eye we examined - especially the retina - appear to be resilient to age-related defects in metabolism, future research will be critical to improve understanding of how the mitochondria in the retina and RPE can influence the broader ocular ecosystem.

## Acknowledgements

The authors would like to thank the animal care teams at the University of Washington that made this study in mice possible. Jeanne Fredrickson helped coordinate logistics of the study. Kelie Gonzalez assisted with genotyping. Finally, the authors thank Susan E. Brockerhoff, Jennifer R. Chao, David *W.* Raible, Andrea E. Wills, Mark A. Kanow, and Christopher C. Farnsworth for thoughtful discussion.

## Funding Information

**NIH/NIA**

- Peter S. Rabinovitch and David J. Marcinek - Mitochondrial Protective Interventions, Aging and Healthspan: 5P01AG001751-36
- Kristine A. Tsantilas T32 - Genetic Approaches to Aging Training Grant T32 AG000057
- Kristine A. Tsantilas T32 - Biological Mechanisms for Healthy Aging Training Grant T32 AG066574
- Mariya T. Sweetwyne K01 - Mitochondrial Protection to Derive Expanded Aged Renal Glomerular Progenitor Cells: K01 AG062757-02

**NIH/NEI**

- James B. Hurley R01 - Determinants of Rod and Cone Response Characteristics: 5R01EY006641
- James B. Hurley R01 - Control of Photoreceptor Metabolism: 5R01EY017863
- Celia M. Bisbach F31 - Understanding the role of cytosolic NADH production in maintaining aerobic glycolysis in the retina: F31EY031165
- Daniel T. Hass T32 - Vision Training Grant: T32EY007031

**NIH/NIDDK**

- Ian R, Sweet - Cell Function Analysis Core of the Diabetes Research Center: P30 DK017047

## Commercial Relationships Disclosures

None to list

## Notes

### Competing Interest Statement

The authors have declared no competing interest.

## References

1. Owsley C. Aging and vision. Vision Res. 2011;51(13):1610–1622. doi:10.1016/j.visres.2010.10.020

2. Klein R, Klein BEK. The prevalence of age-related eye diseases and visual impairment in aging: Current estimates. Investig Ophthalmol Vis Sci. 2013;54(14):ORSF5–ORSF13. doi:10.1167/iovs.13-12789

3. Gu X, Neric NJ, Crabb JS, et al. Age-related changes in the retinal pigment epithelium (RPE). PLoS One. 2012;7(6):e38673. doi:10.1371/journal.pone.0038673

4. Bonilha VL. Age and disease-related structural changes in the retinal pigment epithelium. Clin Ophthalmol. 2008;2(2):413–424. doi:10.2147/opth.s2151

5. Karunadharma PP, Nordgaard CL, Olsen TW, Ferrington DA. Mitochondrial DNA damage as a potential mechanism for Age-Related macular Degeneration. Investig Ophthalmol Vis Sci. 2010;51 (11):5470–5479. doi:10.1167/iovs.10-5429

6. Brunk UT, Terman A. Lipofuscin: Mechanisms of age-related accumulation and influence on cell function. Free Radic Biol Med. 2002;33(5):611–619. doi:10.1016/S0891-5849(02)00959-0

7. Crabb JW. The proteomics of drusen. Cold Spring Harb Perspect Med. 2014;4(7):a017194. doi:10.1101/cshperspect.a017194

8. Chen M, Rajapakse D, Fraczek M, Luo C, Forrester J V., Xu H. Retinal pigment epithelial cell multinucleation in the aging eye - a mechanism to repair damage and maintain homoeostasis. Aging Cell. 2016;15:436–445. doi:10.1111/acel.12447

9. Ambati J, Fowler BJ. Mechanisms of age-related macular degeneration. Neuron. 2012;75(1):26–39. doi:10.1016/j.neuron.2012.06.018

10. Alavi M V. Aging and Vision. In: Bowes Rickman C, LaVail MM, Anderson RE, Grimm C, Hollyfield J, Ash J, eds. Retinal Degenerative Diseases. Vol 854. 1st ed. Springer Nature; 2016:393–399. doi:10.1007/978-3-319-17121-0_52

11. Bhutto I, Lutty G. Understanding age-related macular degeneration (AMD): Relationships between the photoreceptor/retinal pigment epithelium/Bruch’s membrane/choriocapillaris complex. Mol Aspects Med. 2012;33(4):295–317. doi:10.1016/j.mam.2012.04.005.

12. Inana G, Murat C, An W, Yao X, Harris IR, Cao J. RPE phagocytic function declines in age-related macular degeneration and is rescued by human umbilical tissue derived cells. J Transl Med. 2018;16:63. doi:10.1186/s12967-018-1434-6

13. Nag TC, Wadhwa S. Immunolocalisation pattern of complex I-V in ageing human retina: Correlation with mitochondrial ultrastructure. Mitochondrion. 2016;31:20–32. doi:10.1016/j.mito.2016.08.016

14. Wang AL, Lukas TJ, Yuan M, Neufeld AH. Age-related increase in mitochondrial DNA damage and loss of DNA repair capacity in the neural retina. Neurobiol Aging. 2010;31(11):2002–2010. doi:10.1016/j.neurobiolaging.2008.10.019

15. Eells JT. Mitochondrial dysfunction in the aging retina. Biology (Basel). 2019;8(2):31. doi:10.3390/biology8020031

16. Samuel MA, Zhang Y, Meister M, Sanes JR. Age-related alterations in neurons of the mouse retina. J Neurosci. 2011;31(44):16033–16044. doi:10.1523/JNEUROSCI.3580-11.2011

17. Cavallotti C, Artico M, Pescosolido N, Tranquilli Leali FM, Feher J. Age-related changes in the human retina. Can J Ophthalmol. 2004;39:61–68. doi:10.1016/S0008-4182(04)80054-1

18. Wang Y, Grenell A, Zhong F, et al. Metabolic signature of the aging eye in mice. Neurobiol Aging. 2018;71:223–233. doi:10.1016/j.neurobiolaging.2018.07.024

19. Kolesnikov A V., Fan J, Crouch RK, Kefalov VJ. Age-related deterioration of rod vision in mice. J Neurosci. 2010;30(33):11222–11231. doi:10.1523/JNEUROSCI.4239-09.2010

20. Freund PR, Watson J, Gilmour GS, Gaillard F, Sauvé Y. Differential changes in retina function with normal aging in humans. Doc Ophthalmol. 2011;122:177–190. doi:10.1007/s10633-011-9273-2

21. Kergoat H, Kergoat M-J, Justino L. Age-related changes in the flash electroretinogram and oscillatory potentials in individuals age 75 and older. J Am Geriatr Soc. 2001;49:1212–1217. doi:10.1046/j.1532-5415.2001.49239.x

22. Winkler BS. Glycolytic and oxidative metabolism in relation to retinal function. J Gen Physiol. 1981;77(6):667–692. doi:10.1085/jgp.77.6.667

23. Kanow MA, Giarmarco MM, Jankowski C, et al. Biochemical adaptations of the retina and retinal pigment epithelium support a metabolic ecosystem in the vertebrate eye. Elife. 2017;6:e28899. doi:10.1101/143347

24. Haydinger CD, Kittipassorn T, Peet DJ. Power to see—Drivers of aerobic glycolysis in the mammalian retina: A review. Clin Exp Ophthalmol. 2020;48:1057–1071. doi:10.1111/ceo.13833

25. Warburg O. The metabolism of carcinoma cells. J Cancer Res. 1925;9(1):148–163. doi:10.1158/jcr.1925.148

26. Swarup A, Samuels IS, Bell BA, et al. Modulating GLUT1 expression in the RPE decreases glucose levels in the retina: Impact on photoreceptors and Müller glial cells. Am J Physiol Physiol. 2019;316(1):C121–C133.

27. Kurihara T, Westenskow PD, Gantner ML, et al. Hypoxia-induced metabolic stress in retinal pigment epithelial cells is sufficient to induce photoreceptor degeneration. Elife. 2016;5:e14319. doi:10.7554/eLife.14319

28. Zhao C, Yasumura D, Li X, et al. mTOR-mediated dedifferentiation of the retinal pigment epithelium initiates photoreceptor degeneration in mice. J Clin Invest. 2011;121(1):369–383. doi:10.1172/JCI44303

29. López-Otín C, Blasco MA, Partridge L, Serrano M, Kroemer G. The hallmarks of aging. Cell. 2013;153(6):1194–1217. doi:10.1016/j.cell.2013.05.039

30. Du J, Linton JD, Hurley JB. Probing Metabolism in the Intact Retina Using Stable Isotope Tracers Jianhai. Methods Enzymol. 2015;561:149–170. doi:doi:10.1016/bs.mie.2015.04.002

31. Mattapallil MJ, Wawrousek EF, Chan CC, et al. The Rd8 mutation of the Crb1 gene is present in vendor lines of C57BL/6N mice and embryonic stem cells, and confounds ocular induced mutant phenotypes. Invest Ophthalmol Vis Sci. 2012;53(6):2921–2927. doi:10.1167/iovs.12-9662

32. Chang B, Hurd R, Wang J, Nishina P. Survey of Common Eye Diseases in Laboratory Mouse Strains. Investig Ophthalmol Vis Sci. 2013;54(7):4974–4981. doi:10.1167/iovs.13-12289

33. Millard P, Letisse F, Sokol S, Portais JC. IsoCor: correcting MS data in isotope labeling experiments. Bioinformatics. 2012;28(9):1294–1296. doi:10.1093/bioinformatics/bts127

34. Millard P, Delépine B, Guionnet M, Heuillet M, Bellvert F, Létisse F. IsoCor: isotope correction for high-resolution MS labeling experiments. Bioinformatics. 2019;35(21):4484–4487. doi:10.1093/bioinformatics/btz209

35. Sweet IR, Cook DL, Wiseman RW, et al. Dynamic perifusion to maintain and assess isolated pancreatic islets. Diabetes Technol Ther. 2002;4(1):67–76. doi:10.1089/15209150252924111

36. Sweet IR, Khalil G, Wallen AR, et al. Continuous Measurement of Oxygen Consumption by Pancreatic Islets. Diabetes Technol Ther. 2002;4(5):661–672.

37. Yang N-C, Ho W, Chen Y-H, Hu M-L. A convenient one-step extraction of cellular ATP using boiling water for the luciferin-luciferase assay of ATP. Anal Biochem. 2002;306(2):323–327. doi:10.1006/abio.2002.5698

38. Karp NA, Mason J, Beaudet AL, et al. Prevalence of sexual dimorphism in mammalian phenotypic traits. Nat Commun. 2017;8:15475. doi:10.1038/ncomms15475

39. Shaqiri A, Roinishvili M, Grzeczkowski L, et al. Sex-related differences in vision are heterogeneous. Sci Rep. 2018;8:7521. doi:10.1038/s41598-018-25298-8

40. Zetterberg M. Age-related eye disease and gender. Maturitas. 2016;83:19–26. doi:10.1016/j.maturitas.2015.10.005

41. Du M, Mangold CA, Bixler G V., et al. Retinal gene expression responses to aging are sexually divergent. Mol Vis. 2017;23:707–717.

42. Bisbach CM, Hass DT, Robbings BM, et al. Succinate Can Shuttle Reducing Power from the Hypoxic Retina to the O_2_-Rich Pigment Epithelium. Cell Rep. 2020;31(5):107606. doi:10.1016/j.celrep.2020.107606

43. Du J, Yanagida A, Knight K, et al. Reductive carboxylation is a major metabolic pathway in the retinal pigment epithelium. Proc Natl Acad Sci U S A. 2016;113(51):14710–14715. doi:10.1073/pnas.1604572113

44. Jackson Laboratory. Life span as a biomarker. Accessed April 4, 2021. https://www.jax.org/research-and-faculty/research-labs/the-harrison-lab/gerontology/life-span-as-a-biomarker

45. Yuan R, Tsaih S-W, Petkova SB, et al. Aging in inbred strains of mice: Study design and interim report on median lifespans and circulating IGF1 levels. Aging Cell. 2009;8(3):277–287. doi:10.1111/j.1474-9726.2009.00478.x

46. Yuan R, Peters LL, Paigen B. Mice as a mammalian model for research on the genetics of aging. ILAR J. 2011;52(1):4–15. doi:10.1093/ilar.52.1.4

47. Hamberger AC, Chiang GH, Nylén ES, Scheff SW, Cotman CW. Glutamate as a CNS transmitter. I. Evaluation of glucose and glutamine as precursors for the synthesis of preferentially released glutamate. Brain Res. 1979;168(3):513–530. doi:10.1016/0006-8993(79)90306-8

48. Camandola S, Mattson MP. Brain metabolism in health, aging, and neurodegeneration. EMBO J. 2017;36(11):1474–1492. doi:10.15252/embj.201695810

49. Angelin A, Gil-de-Gómez L, Dahiya S, et al. Foxp3 Reprograms T Cell Metabolism to Function in Low-Glucose, High-Lactate Environments. Cell Metab. 2017;25(6):1282–1293. doi:10.1016/j.cmet.2016.12.018

50. Goyal MS, Vlassenko AG, Blazey TM, et al. Loss of Brain Aerobic Glycolysis in Normal Human Aging. Cell Metab. 2017;26(2):353–360. doi:10.1016/j.cmet.2017.07.010

51. Weyand CM, Goronzy JJ. Aging of the immune system: Mechanisms and therapeutic targets. Ann Am Thorac Soc. 2016;13:S422–S428. doi:10.1513/AnnalsATS.201602-095AW

52. Ritterhoff J, Tian R. Metabolismin cardiomyopathy: Every substrate matters. Cardiovasc Res. 2017;113(4):411–421. doi:10.1093/cvr/cvx017

53. Chiao YA, Rabinovitch PS. The aging heart. Cold Spring Harb Perspect Med. 2015;5(9):a025148. doi:10.1101/cshperspect.a025148

54. Rui L. Energy Metabolism in the Liver. Compr Physiol. 2014;4(1):177–197. doi:10.1002/cphy.c130024

55. Garvey SM, Dugle JE, Kennedy AD, et al. Metabolomic profiling reveals severe skeletal muscle group-specific perturbations of metabolism in aged FBN rats. Biogerontology. 2014;15:217–232. doi:10.1007/s10522-014-9492-5

56. Demontis F, Piccirillo R, Goldberg AL, Perrimon N. Mechanisms of skeletal muscle aging: Insights from Drosophila and mammalian models. DMM Dis Model Mech. 2013;6:1339–1352. doi:10.1242/dmm.012559

57. Vitorica J, Satrustegui A, Machado A. Metabolic Implications of Ageing: Changes in Activities of Key Lipogenic and Gluconeogenic Enzymes in the Aged Rat Liver. Enzyme. 1981;26:144–152.

58. Nishikawa T, Bellancce N, Damm A, et al. A switch in the source of ATP production and a loss in capacity to perform glycolysis are hallmarks of hepatocyte failure in advance liver disease. J Hepatol. 2014;60(6):1203–1211. doi:doi:10.1016/j.jhep.2014.02.014

59. Ohlendieck K. Proteomic profiling of fast-to-slow muscle transitions during aging. Front Physiol. 2011; 2(105). doi:10.3389/fphys.2011.00105

60. Wei Y, Jiang H, Shi Y, et al. Age-related alterations in the retinal microvasculature, microcirculation, and microstructure. Investig Ophthalmol Vis Sci. 2017;58(9):3804–3817. doi:10.1167/iovs.17-21460

61. Lin Y, Jiang H, Liu Y, et al. Age-related alterations in retinal tissue perfusion and volumetric vessel density. Investig Ophthalmol Vis Sci. 2019;60(2):685–693. doi:10.1167/iovs.18-25864

62. Orlov N V., Coletta C, van Asten F, et al. Age-related changes of the retinal microvasculature. PLoS One. 2019;14(5):e0215916. doi:10.1371/journal.pone.0215916

63. Bata AM, Fondi K, Szegedi S, et al. Age-Related Decline of Retinal Oxygen Extraction in Healthy Subjects. Investig Ophthalmol Vis Sci. 2019;60(8):3162–3169. doi:10.1167/iovs.18-26234

64. TeSlaa T, Bartman CR, Jankowski CSR, et al. The Source of Glycolytic Intermediates in Mammalian Tissues. Cell Metab. 2021;33(2):367–378. doi:10.1016/j.cmet.2020.12.020

65. Adijanto J, Du J, Moffat C, Seifert EL, Hurley JB, Philp NJ. The retinal pigment epithelium utilizes fatty acids for ketogenesis. J Biol Chem. 2014;289(30):20570–20582. doi:10.1074/jbc.M114.565457

66. Reyes-Reveles J, Dhingra A, Alexander D, Bragin A, Philp NJ, Boesze-Battaglia K. Phagocytosis-dependent ketogenesis in retinal pigment epithelium. J Biol Chem. 2017;292(19):8038–8047. doi:10.1074/jbc.M116.770784

67. Izuta Y, Imada T, Hisamura R, et al. Ketone body 3-hydroxybutyrate mimics calorie restriction via the Nrf2 activator, fumarate, in the retina. Aging Cell. 2017;17(1):e12699. doi:10.1111/acel.12699

68. Yam M, Engel AL, Wang Y, et al. Proline mediates metabolic communication between retinal pigment epithelial cells and the retina. J Biol Chem. 2019;294(26):P10278–10289. doi:10.1074/jbc.RA119.007983

69. Tait SWG, Green DR. Mitochondria and cell signalling. J Cell Sci. 2012;125:807–815. doi:10.1242/jcs.099234

70. Abate M, Festa A, Falco M, et al. Mitochondria as playmakers of apoptosis, autophagy and senescence. Semin Cell Dev Biol. 2020;98:139–153. doi:10.1016/j.semcdb.2019.05.022

71. Hoppins S. The regulation of mitochondrial dynamics. Curr Opin Cell Biol. 2014;29:46–52. doi:10.1016/j.ceb.2014.03.005

72. Bénit P, Letouzé E, Rak M, et al. Unsuspected task for an old team: Succinate, fumarate and other Krebs cycle acids in metabolic remodeling. Biochim Biophys Acta-Bioenerg. 2014;1837(8):1330–1337. doi:10.1016/j.bbabio.2014.03.013

73. Salminen A, Kauppinen A, Hiltunen M, Kaarniranta K. Krebs cycle intermediates regulate DNA and histone methylation: Epigenetic impact on the aging process. Ageing Res Rev. 2014;16:45–65. doi:10.1016/j.arr.2014.05.004

74. Wiley CD, Velarde MC, Lecot P, et al. Mitochondrial dysfunction induces senescence with a distinct secretory phenotype. Cell Metab. 2016;23(2):303–314. doi:10.1016/j.cmet.2015.11.011

75. Bratic A, Larsson NG. The role of mitochondria in aging. J Clin Invest. 2013;123(3):951–957. doi:10.1172/JCI64125

76. Bratic I, Trifunovic A. Mitochondrial energy metabolism and ageing. Biochim Biophys Acta - Bioenerg. 2010; 1797(6-7):961–967. doi:10.1016/j.bbabio.2010.01.004

77. Sun N, Youle RJ, Finkel T. The Mitochondrial Basis of Aging. Mol Cell. 2016;61(5):654–666. doi:10.1016/j.molcel.2016.01.028

