## Supplemental Material for "Metabolic function in aging retina and retinal pigment epithelium remains robust despite vision loss"

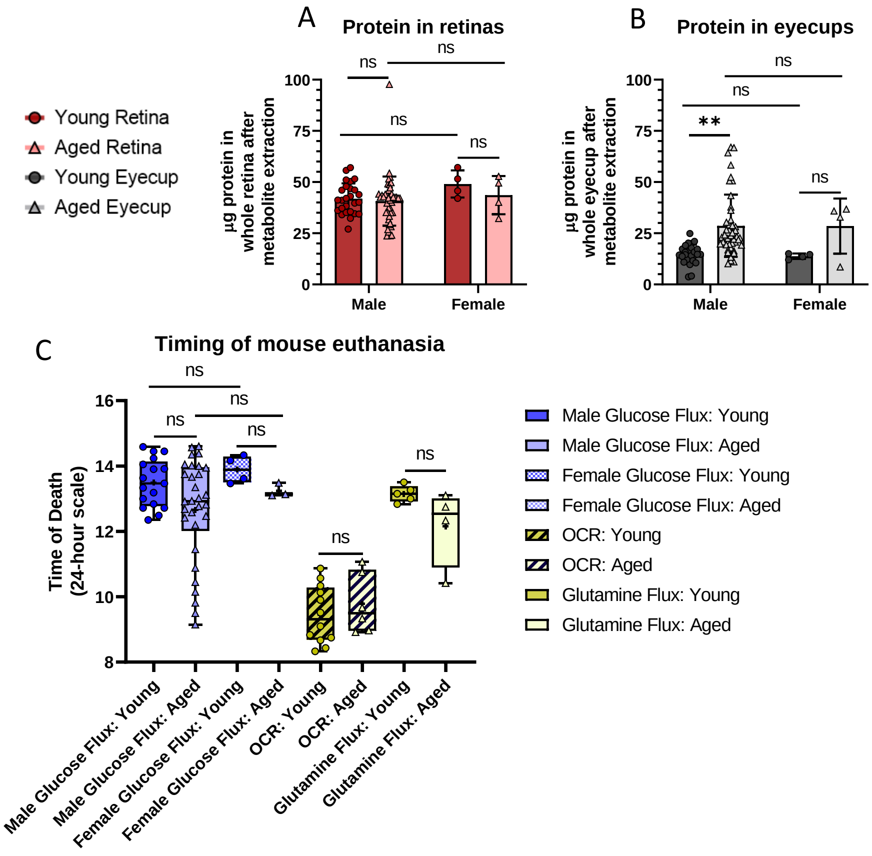


**Supplementary Figure 1:** Sample collection time and quantity of tissue were considered during this study. We observed no significant difference in the protein content between young and aged retinas (A) of either sex in our glucose timecourse. A group of eyecup samples trended significantly higher (B) and increased the average, but this was found to correlate with the individual who collected the samples. Differences in tissue quantity were controlled for in the study by normalizing all values to total protein in the tissue. Values are plotted as the mean ± standard deviation. The normality of data was determined using the Shapiro-Wilk test. Age and sex-related differences were tested (* = $p < 0.05$, ** = $p < 0.0001$) using the Kruskal-Wallis and Dunn's multiple comparison tests. The time of death for all animals used in the flux and OCR experiments (C) except for a single aged female mouse, whose time of death was not noted but was known to occur between the hours plotted. We did not observe any clear differences related to the time of day the sample was collected. Boxes represent the median, minimum, and maximum. Means are marked with a cross (+). The normality of data was determined using the Shapiro-Wilk test. The age and sex-related differences in glucose flux were tested using the Kruskal-Wallis and Dunn's multiple comparison tests. Age-related differences in glutamine flux and OCR were tested using unpaired t-tests.

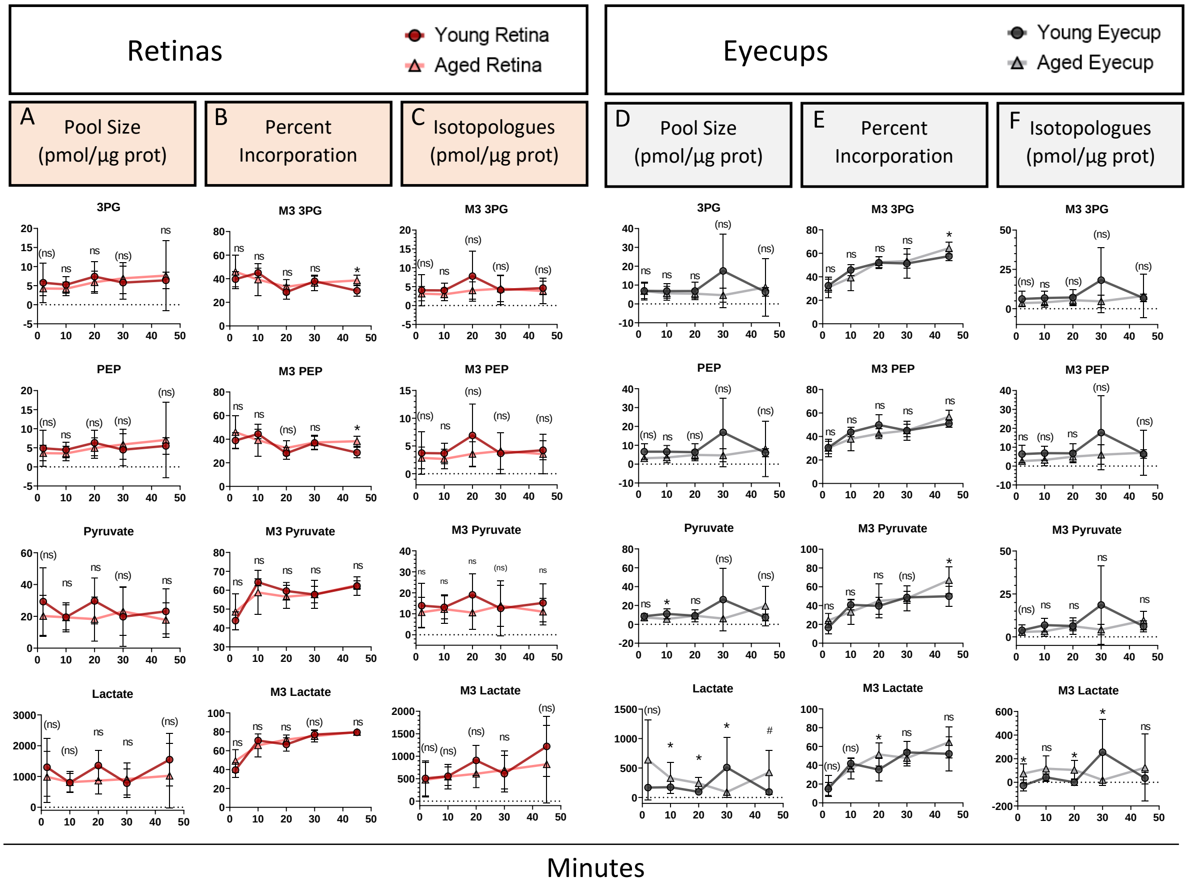


**Supplementary Figure 2:** Individually plotted glycolytic intermediates from the glucose time course in retinas and eyecups. The pool size (A), percent $^{13}C$ incorporation (B), and isotopologues (C) in the retina are listed for select glycolytic intermediates. In eyecups for the same glycolytic intermediates, the pool size (D), percent $^{13}C$ incorporation (E), and isotopologues (F) are shown. Values shown are the mean ± standard deviation. Sample size = 4-9 depending on the age, tissue, and timepoint. Note that these graphs and sample sizes consider two outliers that were removed by Grubb's test (alpha = 0.05, $p < 0.05$): one young retina at 20 minutes and one aged eyecup at 30 minutes. Both were more than 10-fold higher than other tissues at the same timepoints. Normality of data was determined using the Shapiro-Wilk test and p-values were calculated for age-related comparisons using unpaired t-tests (* = $p < 0.05$) or Mann-Whitney tests (# = $p < 0.05$, marker of non-significance enclosed in parentheses). Error bars represent the standard deviation.

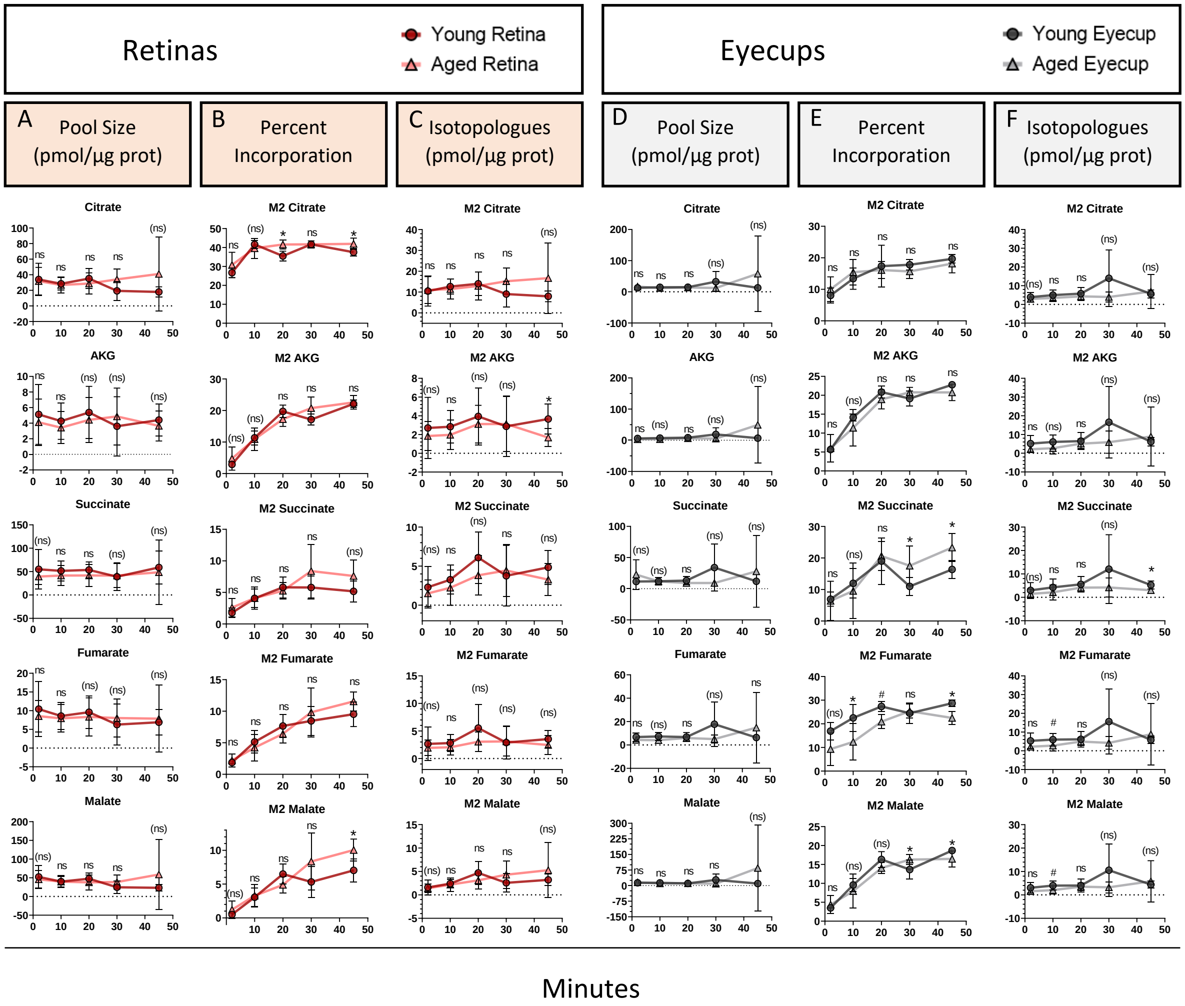


**Supplementary Figure 3:** Individually plotted Krebs cycle intermediates from the glucose time course in retinas and eyecups. The pool size (A), percent $^{13}C$ incorporation (B), and isotopologues (C) in the retina are listed for select Krebs cycle intermediates. In eyecups for the same intermediates, the pool size (D), percent $^{13}C$ incorporation (E), and isotopologues (F) are shown. Values shown are the mean ± standard deviation. Sample size = 4-9 depending on the age, tissue, and timepoint. Note that these graphs and sample sizes consider two outliers that were removed by Grubb's test (alpha = 0.05, $p < 0.05$): one young retina at 20 minutes and one aged eyecup at 30 minutes. Both were more than 10-fold higher than other tissues at the same timepoints. Normality of data was determined using the Shapiro-Wilk test and p-values were calculated for age-related comparisons using unpaired t-tests (* = $p < 0.05$) or Mann-Whitney tests (# = $p < 0.05$, marker of non-significance enclosed in parentheses). Error bars represent the standard deviation.

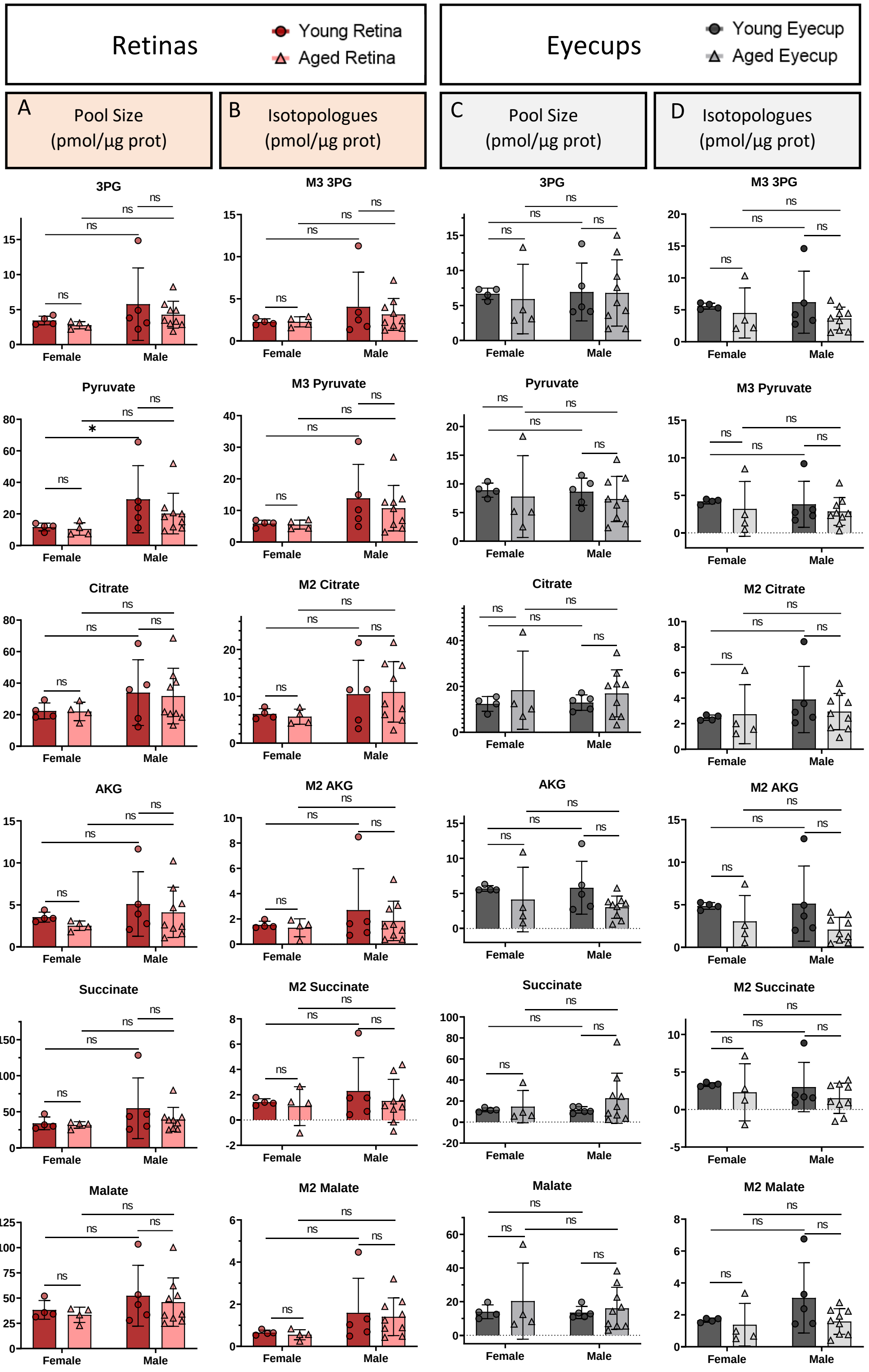


**Supplementary Figure 4:** Sex differences between males (26 months) and females (23-25 months) were examined by individually plotting glycolytic and Krebs cycle intermediates measured in young and aged mice after a 2 minute incubation in 5 mM U-$^{13}$C-glucose. Note that the male values are the same 2 minute samples as shown in Figure 3 and Supplementary Figures 3 and 4. The pool size (A) and isotopologues (B) in the young and aged retina of both sexes are listed for select glycolytic and Krebs cycle intermediates. In eyecups for the same metabolites, the pool size (C) and isotopologues (D) are shown. Values shown are the mean ± standard deviation. Sample size = 4-9 depending on age and sex. Normality of data was determined using the Shapiro-Wilk test and changes associated with aging and sex were examined using Kruskal-Wallis and Dunn's multiple comparison tests.

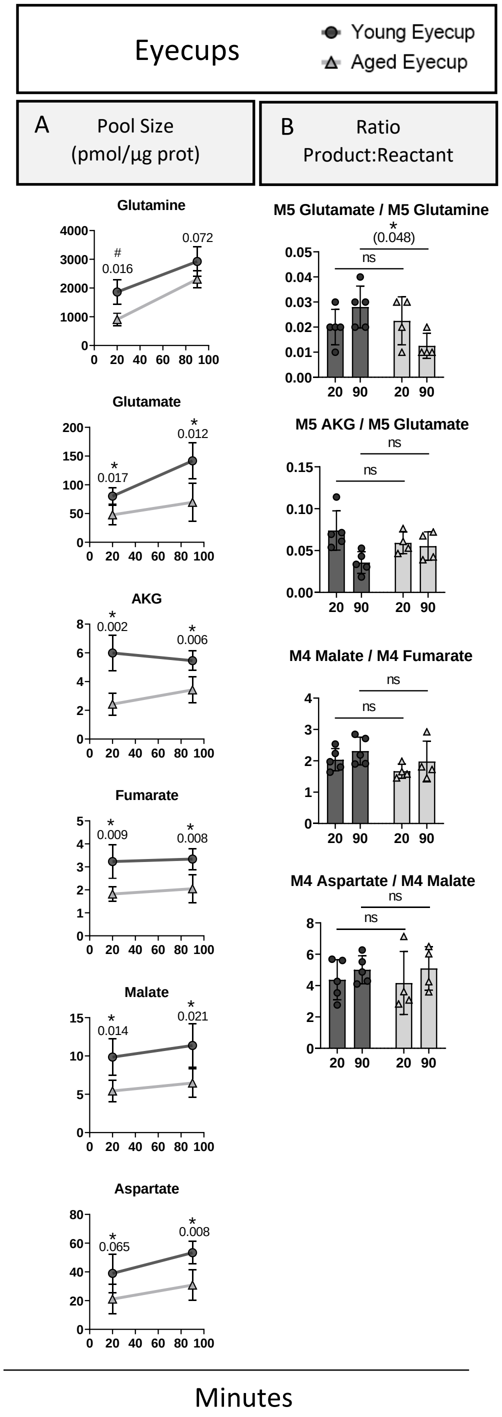


**Supplementary Figure 5:** Intermediates entering and within the Krebs cycle from a glutamine time course (20 and 90 minutes) in eyecups. Pool sizes (A) are generally lower in aged eyecups, while product:reactant ratios (B) were found to have minimal significant changes. Only the ratio of M5 glutamate/M5 glutamine decreased slightly in aged eyecups. Although succinate isotopologues were searched for in these experiments, they were not reliably above the limit of detection in eyecups, thus we could not determine any age-related changes involving succinate in eyecups. Values shown are the mean ± standard deviation. Sample size is 5 for young and 4 for aged at both timepoints. Normality of data was determined using the Shapiro-Wilk test and p-values were calculated using unpaired t-tests (* = $p < 0.05$) or Mann-Whitney tests (# = $p < 0.05$, marker of non-significance enclosed in parentheses).

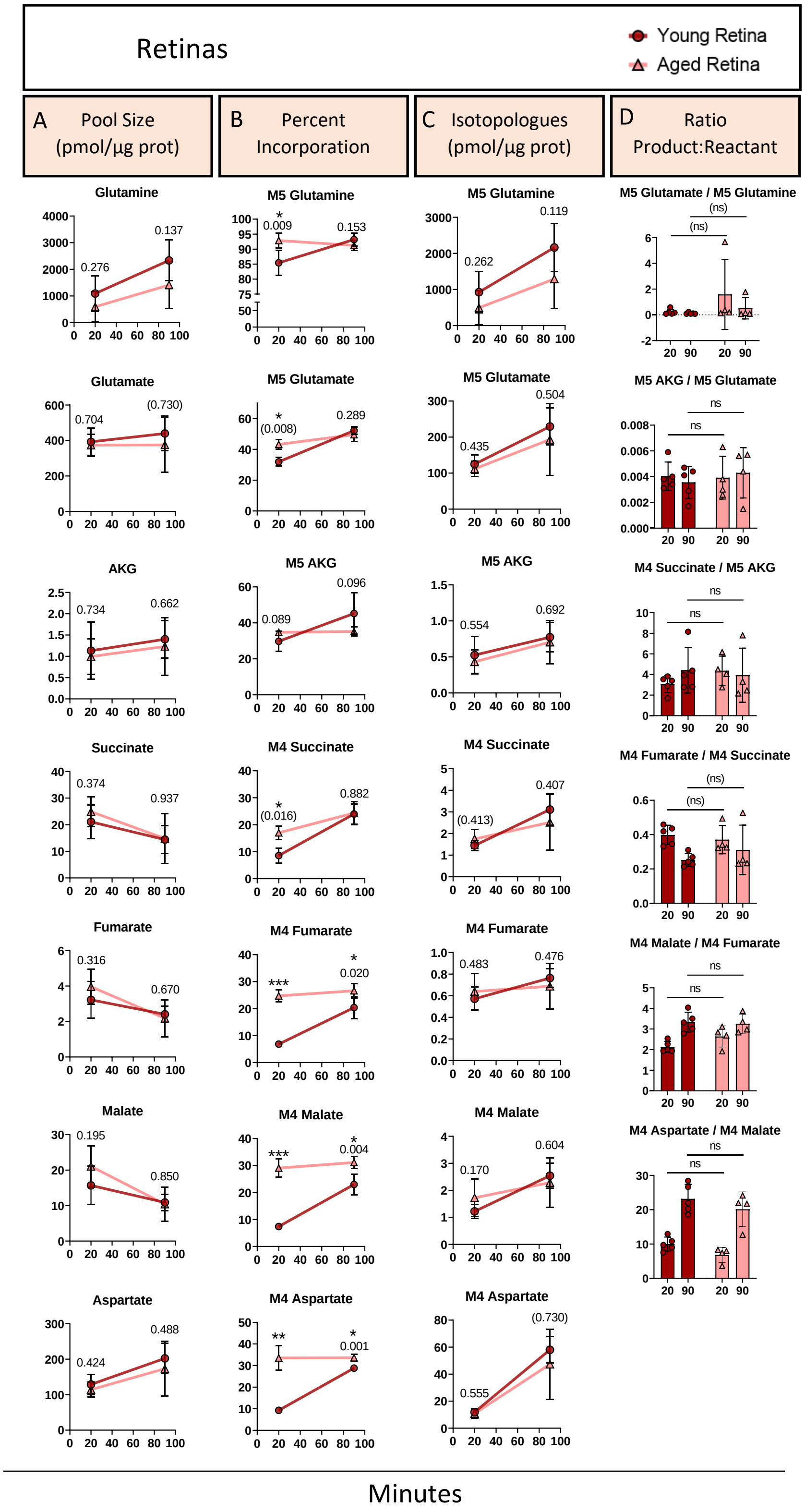


**Supplementary Figure 6:** Examining intermediates entering and within the Krebs cycle from a glutamine time course (20 and 90 minutes) in retinas. The pool size (A) is unchanged. The percent $^{13}C$ incorporation (B) remains unchanged in aged retinas at both times, while it is consistently lower in young retinas at 20 minutes for all intermediates (except for AKG) but has matched the aged by 90 minutes. The quantity of labeled isotopologues (C), and the product:reactant ratios (D) in the retina show no significant changes. Values shown are the mean ± standard deviation. Sample size is 5 for young and 4 for aged at both timepoints. Normality of data was determined using the Shapiro-Wilk test and p-values were calculated using unpaired t-tests (* = $p < 0.05$) or Mann-Whitney tests (# = $p < 0.05$, marker of non-significance enclosed in parentheses).

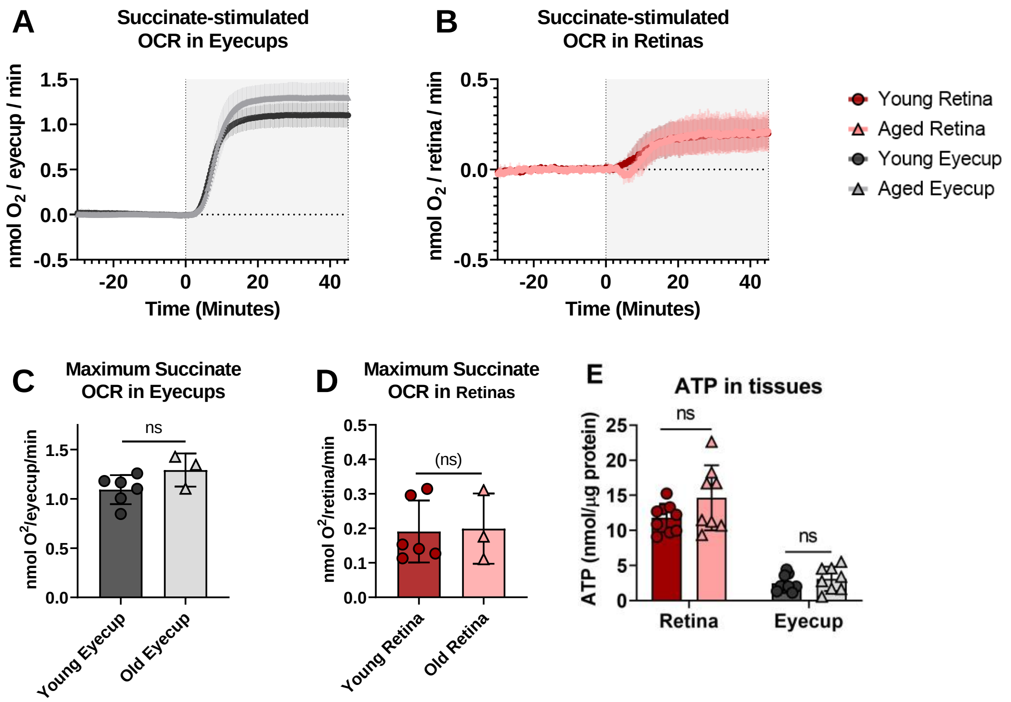


**Supplementary Figure 7:** Oxygen consumption rate was measured in terms of nmol $O_2$ per retina or eyecup per minute using a continuous perifusion system. The values were normalized by subtracting the baseline consumption in 5 mM glucose for eyecups (A) and retina (B). The basal oxygen consumption values (not normalized) are listed in Supplementary Table 1. There was a modest increase in aged eyecup OCR in response to succinate (Shaded area between 0-45 minutes), although it does not reach statistical significance. There was no discernable change with age in retinas (D). We observed no substantial changes in steady-state ATP levels with age when measured using the Molecular Probes® ATP Determination Kit (E). Normality of data was determined using the Shapiro-Wilk test and p-values were calculated) for age-related comparisons using unpaired t-tests (* = $p < 0.05$) or Mann-Whitney tests (# = $p < 0.05$, marker of non-significance enclosed in parentheses). Error bars represent the standard deviation.

**Supplementary Table 1:** Average *ex vivo* $O_2$ consumption in young and aged retina and eyecups

| Tissue | Age | n | Basal $O_2$ consumption* (nmol $O_2$/min) ± S.D. | Max $O_2$ consumption with succinate† (nmol $O_2$/min) ± S.D. |
|---|---|---|---|---|
| Retina | Young | 6 | 1.90 ± 0.4 | 2.04 ± 0.4 |
| | Aged | 3 | 2.14 ± 0.4 | 2.34 ± 0.4 |
| Eyecup | Young | 6 | 0.52 ± 0.3 | 1.62 ± 0.3 |
| | Aged | 3 | 0.56 ± 0.2 | 1.85 ± 0.2 |

* Basal $O_2$ measured between -30 and 0 minutes on the normalized graphs

† Maximum $O_2$ with succinate was measured between 20 and 45 minutes on the normalized graphs

**Supplementary Table 2:** Metabolite standards for method development and normalization.

| Metabolite | Product number | Retention Time 1* (mins) | Retention Time 2† (mins) | Derivatized Mass | # TBDMS | # MeOx | Target Fragment | Target Ions | Qualifier Fragment | Qualifier Ion |
|---|---|---|---|---|---|---|---|---|---|---|
| 3-PG | P7127 | 37.2 | 36.27 | 642 | 4 | 0 | M-57 | 585-588 | M-159 | 483 |
| α-ketoglutarate | 75890 | 26.99 | 25.97 | 403 | 2 | 1 | M-57 | 346-351 | M-15 | 388 |
| Alanine | A7627 | 16.86 | 15.93 | 317 | 2 | 0 | M-57 | 260-263 | M-85 | 232 |
| Aspartate | A6558 | 29.66 | 28.56 | 475 | 3 | 0 | M-57 | 418-422 | M-85 | 316 |
| Citrate | S4641 | 37.1 | 36.12 | 648 | 4 | 0 | M-57 | 591-597 | M-189 | 459 |
| DHAP | D7137 | 32.84 | 32.15 | 541 | 3 | 1 | M-57 | 484-487 | M-85 | 526 |
| Fumarate | F1506 | 22.8 | 21.53 | 344 | 2 | 0 | M-57 | 287-291 | M-15 | 329 |
| GAP | G5251 | 32.45 | 31.75 | 541 | 3 | 1 | M-57 | 484-487 | M-85 | 456 |
| Glutamate | G8415 | 31.71 | 30.6 | 489 | 3 | 0 | M-57 | 432-437 | M-159 | 330 |
| Glutamine | G3126 | 34.23 | 33.06 | 488 | 3 | 0 | M-57 | 431-436 | M-159 | 329 |
| Lactate | L7022 | 15.51 | 12.8 | 318 | 2 | 0 | M-57 | 261-264 | M-85 | 233 |
| Malate | M1000 | 28.82 | 27.83 | 476 | 3 | 0 | M-57 | 419-423 | M-15 | 461 |
| Methionine | M5308 | 26.14 | 25.09 | 377 | 2 | 0 | M-57 | 320-325 | M-85 | 292 |
| Methylsuccinate | M81209 | 22.17 | 21.09 | 360 | 2 | 0 | M-57 | 303 | M-15 | 345 |
| Norleucine | N6877 | 21.51 | 20.66 | 359 | 2 | 0 | M-57 | 302-308 | M-85 | 274 |
| Norvaline | N7627 | 19.88 | 18.95 | 345 | 2 | 0 | M-57 | 288-293 | M-85 | 260 |
| PEP | P7127 | 30.85 | 29.9 | 510 | 3 | 0 | M-57 | 453-456 | M-15 | 495 |
| Pyruvate | P4562 | 9.58 | 8.87 | 231 | 1 | 1 | M-57 | 174-177 | M-131 | 100 |
| Serine | S4500 | 26.49 | 25.56 | 447 | 3 | 0 | M-57 | 390-393 | M-159 | 288 |
| Succinate | 14160 | 22.04 | 20.89 | 346 | 2 | 0 | M-57 | 289-293 | M-15 | 331 |

* In method 1, a DB-5MS column with a flow rate of 0.8 mL/min was used. Column length was 25 m with an inner diameter of 200 µm, and a 0.33 µm nonpolar phenyl arylene polymer film.

† In method 2, a HP-5MS with a flow rate of 1 mL/min was used. Column length was 30 m with an inner diameter of 250 µm, and a 0.25 µm 5% phenyl methyl silox film.
